# Reproductive restraint to avoid the costs of reproductive conflict in a cooperatively breeding mammal

**DOI:** 10.1101/2024.03.12.584411

**Authors:** Graham Birch, Hazel J. Nichols, Francis Mwanguhya, Michael A. Cant, Jonathan D. Blount

**Affiliations:** Centre for Ecology & Conservation, Faculty of Environment, Science & Economy, University of Exeter, Penryn Campus, Cornwall, TR10 9FE, UK

**Keywords:** Life-history, reproductive conflict, cooperative breeder, restraint, reproductive costs, male

## Abstract

The costs of reproductive conflict can be an important factor shaping the evolution of life histories in animal societies. These costs may change as individuals age and grow, and with within-group competition. Social costs of reproductive conflict have been invoked to explain why females might gain from delaying maturity or ceasing reproduction midway through life, but not in males. Here we analyze more than 20 years of data to understand how individual male banded mongooses adjust their reproductive activity in response to the costs of reproductive conflict. In banded mongooses groups multiple female breeders enter oestrus synchronously and males compete to mate guard the females. The oldest and heaviest males in the group gained the greatest share of paternity, while younger and lighter males obtained little or no paternity. Those lighter males that are reproductively active paid disproportionate survival costs. Our results suggest that by delaying or suppressing reproduction early in life, males may reduce their mortality. Males may also avoid inflicting costs on their male kin by queuing. These results from an egalitarian cooperative breeding mammal may provide a rare example of voluntary reproductive restraint, although we cannot discount reproductive suppression by rival males.

## Introduction

The assumption that reproduction involves costs to future fecundity and/or survival has formed the bedrock of life-history theory since its inception [1–8]. Due to finite resources, individuals are expected to balance the costs and benefits of reproduction across their lifespan to maximise their lifetime reproductive success [5–7]. These costs and benefits can vary according to an individual’s condition. Individuals in good condition might escape or reduce life history tradeoffs as they have more metabolic resources to put towards both reproductive effort and somatic maintenance [9,10]. Condition changes as individuals age, with older and heavier individuals having more resources to put towards reproductive effort, making them typically the most fecund or successful competitors in a population [11,12]. Males that engage at a younger age may gain more opportunities for reproductive success but should face higher costs of reproductive activity if they are in poorer condition. For example, reproduction in early adulthood is associated with accelerated senescence compared to later onset of reproduction in red deer (*Cervus elaphus*) [13,14]. Consequently, reproductive life-histories should be optimised according to this change in condition as individuals develop and age to maximise lifetime reproductive success.

The costs and benefits of reproduction are also shaped by changing environment. An unfavourable environment may favour a delay or pause in reproductive activity. For example, African striped mice (*Rhabdomys pumilio*) delay dispersal under high population densities [15]. In social groups the costs and benefits of reproduction vary with the social environment, such as the number and condition of helpers, mating partners, and rivals in the group. For example, offspring have worse survival outcomes in groups with only genetically incompatible or low quality mating partners, or in cooperative breeding groups with few helpers [16–18]. Such conditions could favour reduced reproductive investment, particularly if the social environment is likely to be more favourable later in life. The intensity of reproductive conflict, defined in this study as pre-copulatory competition over mating partners, is also likely to vary across the lifespan. The relative condition and resource holding potential (RHP) of an individual compared to same-sex rivals within the group should determine the social costs of reproductive conflict for that individual and its likelihood of gaining reproductive fitness [19].

Here we examine how reproductive life-histories are adjusted in response to variation in the costs and benefits of reproductive conflict. Reproductive conflict has been proposed to act as a moulding force on life-histories. In early adulthood, social costs imposed by higher RHP group members can supress subordinate reproduction through threats and violence [18,20–27]. The degree of suppression varies across social groups, from despotic high skew groups where only a single dominant pair breeds to more egalitarian lower skew groups where reproduction is more evenly distributed among members [19,28]. In many high skew cooperative breeding groups, conflict over social rank or reproductive access can escalate to the point of evicting competitors [29–32]. Such evictions are exceptionally costly for subordinates who lose the help and protection of the group for themselves and their potential offspring [33], resulting in a high failure rate for establishment of new groups [16,34]. To avoid these social costs, young subordinates can delay their reproductive life-history by remaining inactive waiting in a queue to inherit future reproductive roles [29,34,35]. They may appease dominant individuals through submissive behaviours [36,37], restricting growth to appear less of a threat [38,39], and ‘paying to stay’ through cooperative behaviour such as offspring care [36]. The social costs of suppression can force subordinates to delay their reproductive activity until they gain the RHP necessary to challenge for dominant reproductive positions in social groups.

Kin selected costs of social conflict can also favour delays in reproductive activity. In cooperatively breeding groups, group members are often closely related to each other [19,40,41], although intragroup relatedness can vary substantially, particularly in groups with multiple breeders [42]. Individuals can benefit from staying in their group even if they do not ultimately secure reproductive roles themselves, as they can enhance their inclusive fitness by helping relatives [43]. These indirect benefits may serve to stabilise reproductive queues in cooperative breeding groups [44,45]. In the later stages of life an individual’s relatedness to the group may increase as more offspring, including those of close relatives, are produced and mature [46,47]. Reproduction may increasingly come with kin-selected costs as individuals age, when the offspring of overlapping generations compete with each other for resources and care provided by the group [47–49]. The condition of individuals may also decline as they begin to senesce, reducing their capacity to take on the costs of reproductive investment[50]. The inclusive fitness benefits of helping may therefore start to outweigh the net benefits of reproduction at senescent ages, which can select for reproductive restraint. For example, menopause has been proposed to evolve as form of reproductive restraint in grandmothers. In killer whales (*Orcinus orca*) [51] and some human societies [52] there is evidence that older females can gain from permanently ceasing reproduction midway through life (i.e. to undergo menopause) to allow their younger female relatives to reproduce [51]. By reducing individual reproductive costs at old age, grandmothers can also extend their post-reproductive lifespan to continue assisting their relatives [46]. The moulding of social life histories to reduce reproductive conflict within social groups has been termed ‘life history segregation’ [19].

Examples demonstrating how life-histories adjust to the social and kin selected costs of reproductive conflict have predominantly come from females. Much less is known about adjustments to male reproductive life-histories in social groups, despite the significant costs associated with reproductive competition among males [53–57]. This knowledge gap may be explained by the less tractable nature of male reproductive life-histories in social groups. It is challenging to collect accurate long-term data on the reproductive history of males. Copulations are often difficult to observe compared to the relatively conspicuous changes in the bodies and behaviours of reproducing females, and it can be impossible to confirm paternity without the availability of genetic data. Additionally, many males are obligate dispersers so it may not be necessary to adjust their reproductive life-histories to conflict with kin. Moreover, large dispersal distances provide difficulties identifying the past life-history of males that form or join non-natal groups. Where these problems can be overcome, how reproductive conflict alters the reproductive life-histories of males deserves wider assessment.

We use a comprehensive dataset spanning more than two decades, comprising behavioural, genetic, and life-history data to explore how reproductive activity adjusts to the costs and benefits of reproductive conflict in male banded mongooses. Banded mongooses live in large groups of up to 60 individuals, although groups of 10-30 are more typical (median = 18 adults, interquartile range = 9.25) [58,59]. The most common cause of adult mortality in our study population is predation, with disease or death attributed to senescence being less common [59]. Both sexes reach sexual maturity at around 1 year of age [60]. Banded mongooses are an intermediate skew cooperative breeder with reproductive opportunities spread across a ‘core’ of breeding adults (1–5 females and 3–7 males) that reproduce 3–4 times per year [59], while other male group members remain reproductively inactive. Due to the higher survival rate of males and their tendency to stay in their natal groups, most groups have heavily male biased sex ratios [58]. This skewed sex ratio sets the stage for reproductive conflict over a limited number of breeding females.

All adult females in banded mongoose groups enter oestrus over a period of 7-10 days, preventing their monopolisation by a single dominant male [58]. On average, older females enter oestrus first, followed by younger females a few days later [58]. The period from the first to last signs of oestrus within a group is labelled an ‘oestrus event’. Reproductive males guard individual females by following them closely and aggressively defending them from other males [58]. Guarding males remain active throughout the oestrus event, noticeably reducing foraging effort compared to inactive males. Older males tend to guard older, more reproductively succesful females, despite efforts by females to escape their guards [58].

Here we investigated how male banded mongooses adjust their reproductive activity between oestrus events throughout their lives. Specifically, we employed state-transition models to firstly ask whether there is evidence that males delay the onset of reproductive activity when they are at an RHP (weight and age) disadvantage compared to their competitors. Second, we asked how males decline in reproductive activity as they senesce. We expected RHP to decrease at the oldest age classes, in line with previously identified weight declines in aged banded mongooses [61], and in other taxa [62]. We further assessed the degree to which reproductive fitness is dominated by the highest RHP males in the group, and whether there is evidence that fitness declines rapidly in old age. We also asked whether males suffer energetic costs in the form of weight loss as a result of engaging in reproductive activity. Finally, we asked whether any social and energetic costs of reproductive activity contribute to mortality, and if low RHP or condition males are at a higher risk of death. We predicted that males of lower RHP relative to rivals should experience higher mortality costs when reproductively active. We also predicted higher mortality costs of reproductive activity at very old ages. Simultaneous assessment of patterns of reproductive activity and mortality allowed us to judge if the costs of reproductive activity shape the reproductive life-histories of male banded mongooses.

## Methods

### Study population

We collected data from a banded mongoose population living on the Mweya Peninsula, Queen Elizabeth National Park, Uganda (0°12′S, 29°54′E) between April of 2003 and February 2021. For climate and habitat details see [59]. Each individual was given a unique fur shave patterns on the small area of their back for identification. The history of each individual and group membership is known through life-history data collection ongoing since 1995 [59].

### Reproductive behaviour data collection

Groups were visited at least every 3 days to collect life-history data. Where signs of oestrus were found, groups were visited every day until reproductive behaviour ceased. Each day during oestrus every breeding female received independent observation (focals) lasting 20 minutes. During focals, any males that guarded the focal female were noted. Guards were identified as male that followed the focal female throughout the 20 minutes within a 5 metre distance. Focals took place during peak foraging periods in the morning, and were paused where view of the focal female was obscured, or during group alarms. Groups continued to be visited every day to collect focals until reproductive behaviour ceased. Data on males younger than subadults (less than 180 days of age) was discounted in all analyses.

Most oestrus events (events spanning the start of oestrus until reproductive behaviours cease) had multiple days of data collection (n= 379, mean:2.997, IQR:3). On each day males were noted as a guard or pesterer according to their behaviour towards females, and as a subordinate if they showed no interest. For the purposes of examining behaviour transitions between oestrus events, male reproductive behaviour from all data collection days was summarised into one state for each oestrus event. If males guarded on 50% or more data collection days males were defined as a guard, whilst males were assigned as subordinates if they were inactive for the whole oestrus event. Males that did not guard on 50% or more days, but instead pestered on more than 50% of data collection days, were defined as a pesterer.

### Life-history and weight data collection

All males in this analysis were followed from birth to death and are therefore of known age. The demography of groups is stratified around surviving siblings of shared litters, meaning many males share age ranks. In our analysis shared age ranks were assigned to the minimum rank and subsequent younger males’ ranks left a gap, for example if there were 3 age rank 2 males the next oldest was assigned age rank 5.

Individual body weights have been collected for the population since 2000. Weight collection frequency varies between individuals and groups depending on their habituation. To account for this variation, weights were averaged over a shared time period. For the state transition and paternity analysis, weights were averaged for the 60 days before and after a given oestrus event (oestrus weight). We then calculated relative weights compared to the group average (group centred weights: weight – mean group weight). 7.0% (209/2999; state transition analysis) and 6.5% (90/1386; paternity analysis) of oestrus weights could not be calculated with data available. Imputation has been used in previous studies on long term populations where missing data can occur [63,64], including this same population of banded mongooses [61]. Using the full history of weight collection for each individual, and the correlation between age and weight in banded mongooses, these missing weights were inputted (see supplementary for more detail).

For weight loss models, prior and post oestrus weights were extracted separately without imputation. Prior oestrus weights were averaged over the 60 days prior to each oestrus event and the post oestrus weight was the nearest single weight recorded after an oestrus event ceased (post-oestrus weight). A preliminary model was run to detect an effect of time to post-oestrus weight collection on weight loss, of which no significant effect was found when restricting sampling to one week post-oestrus. Males with missing weights were simply not used.

### Pedigree data

A genetic pedigree has been collected since 2003 [see references for how the pedigree is obtained;11,40]). A banded mongoose’s gestation period is around 9 weeks [pooled data from 58,65]. 105 group litters (402 offspring) could be connected to 180 sires in oestrus events approxiamtely 9 weeks prior to the litter’s birth date (ended 59 ±15 days before).

### Statistical models

All models were fit using Bayesian inference (JAGS MCMC) in R[66]. To improve model convergence, numeric covariates with a range below 0 and above 1 were standardised. Multicolinearity was checked using the ggmcmc package[67]. Chain convergence was checked using rhat values from the JAGS model output, with all models showing convergence of chains for each fitted parameter (R<1.1). Convergence was also checked using traceplots by eye (ggmcmc).

### Mortality and state transition modelling

In this analysis we aimed to focus on active mate guarding and inactive subordinate roles. Secondary reproductive roles (sneakers) are present but were found to be rare for all values of age and weight (<0.10 probability), so for simplification, model outputs involving transitions from or too secondary reproductive roles have been omitted. Males must have been involved in at-least two oestrus events for transition probabilities to be calculated, so males that only lived through a singular observed oestrus event are not included in these models.

The transition probability of males from one reproductive status to the next, and state specific probability of dying before the next oestrus event, was modelled in relation to male age, age rank, and group centred weight using JAGS MCMC [66,68]. This model was based on 2999 state transitions in total. The model used state matrix **z** with element **z***_i,t_* for the state of male *i* at oestrus event *t. i* comprises the 320 males that were followed from birth to death in the study population, that lived through at-least two oestrus events. *t* comprises each oestrus event in order a given male has been present for throughout their life. *A* state-transition matrix **Ω** was set with four dimensions, previous reproductive state *n*, new state *m*, male *i,* and oestrus event *t*. The state process **ω***_n,m,I,t_* represents the probability that a male at reproductive state *n* at oestrus event *t*, will be in state *m* at the next oestrus event *t+1*. The probability of leaving any reproductive status and dying before the next oestrus event was defined as 1 minus the reproductive status related survival probability. Once a male had died its probability of remaining dead was defined as 1. Since we have complete records of group composition during oestrus events, and each male *i* had reproductive state data for each oestrus event *t* they lived through, an observation matrix was not required (always observed). Using this state-transition matrix **Ω** mortality (Iterations=50000, Thinning interval =100, burn in=5000, Chains=3) and reproductive state transitions (Iterations=20000, Thinning interval =100, burn in=2000, Chains=3) were modelled separately.

To allow for the assessment of a separate senescence effect while avoiding problems with multicollinearity, state transitions and mortality models were run twice-first with age rank and second with quadratic age. Group centred weight was included in each age model variation. Age rank and group centred weight (r=0.44), and age and group centred weight (r=0.37), were moderately correlated and did not produce problems with model convergence. Indeed, running separate models for moderately correlated variables may lead to biasing exaggerating their significance [69]. Interactions between weight and age were, however, not considered since interactions between moderately correlated variables produce significant multicolinerarity problems [69].

To control for common group membership during oestrus events and repeated sampling of the same males, random effects for oestrus event ID (n=375), group ID (n=20), and male ID (n=320) were fitted for state transition models. To control for probability of death increasing with time, mortality transition models were also fitted with time to next oestrus event. To remove cases where excessive time had passed before death since the last oestrus event, data were truncated so that males that died more than a year from the last oestrus event were removed from analysis (random effects fitted: oestrus event ID (n=371), historical group ID (n=20), and male ID (n=280)). Initially mortality analysis included time interactions with all fixed effects, but these were later removed when none proved credible.

### Fitness models

The number of offspring sired by each male in a given oestrus event was regressed against group centered weight with a binomial error structure (iterations=20000, thinning interval =100, burn in=2000, chains=3), with total offspring sired in an oestrus event used as the maximum number of successes. Mirroring the transition and mortality models, two model variations were run, one including age rank and the other quadratic age. To control for common group membership and repeated sampling on individuals, random effects for oestrus event id (n=105), historical group ID (n=11), and male ID (n=388) were fitted.

### Weight loss models

554 weight changes from subordinates, and 265 weight changes from guards were included. To control for common group membership during oestrus events and repeated sampling on the same males, random effects for oestrus event ID (n=118), historical group ID (n=20), and male ID (n=259) were fitted. Percentage weight loss for each individual over an oestrus event was normally distributed, and as such was regressed with behavioural state (subordinate vs guard) using a Gaussian distribution using the JAGMS MCMC engine (iterations=20000, thinning interval =100, burn in=2000, chains=3). Initially we wanted to assess whether age had an effect on weight loss for guards compared to subordinates. However age, or age in interaction with state, was not fitted due to issues with multicollinearity with state.

### Model output processing

For all models, significance table outputs were generated using the MCMCvis package[70], and model diagnostic plots using ggmcmc. Plotted probabilities were simulated from the posterior distribution extracted using the JAGSUI package[71] (Figures 1, 2, 3,S3).

### Ethics

Prior approval of all work was received from Uganda Wildlife Authority (UWA) and Uganda National Council for Science and Technology (UNCST). The Ethical Review Committee of the University of XXXXX approved all research activites.

### Results

On average, subordinates were more likely to stay as a subordinate in the next oestrus event (figure 1ai; mean=0.72, hci=0.756, lci=0.644) than to gain a guarding role (mean=0.2,, hci=0.28, lci=0.16), while transition probabilities from a guarding role were similar (figure 1bi; guard to subordinate-mean=0.46, hci=0.523, lci=0.386; stay guard – mean=0.41, hci=0.5, lci=0.34).

**Figure 1:**
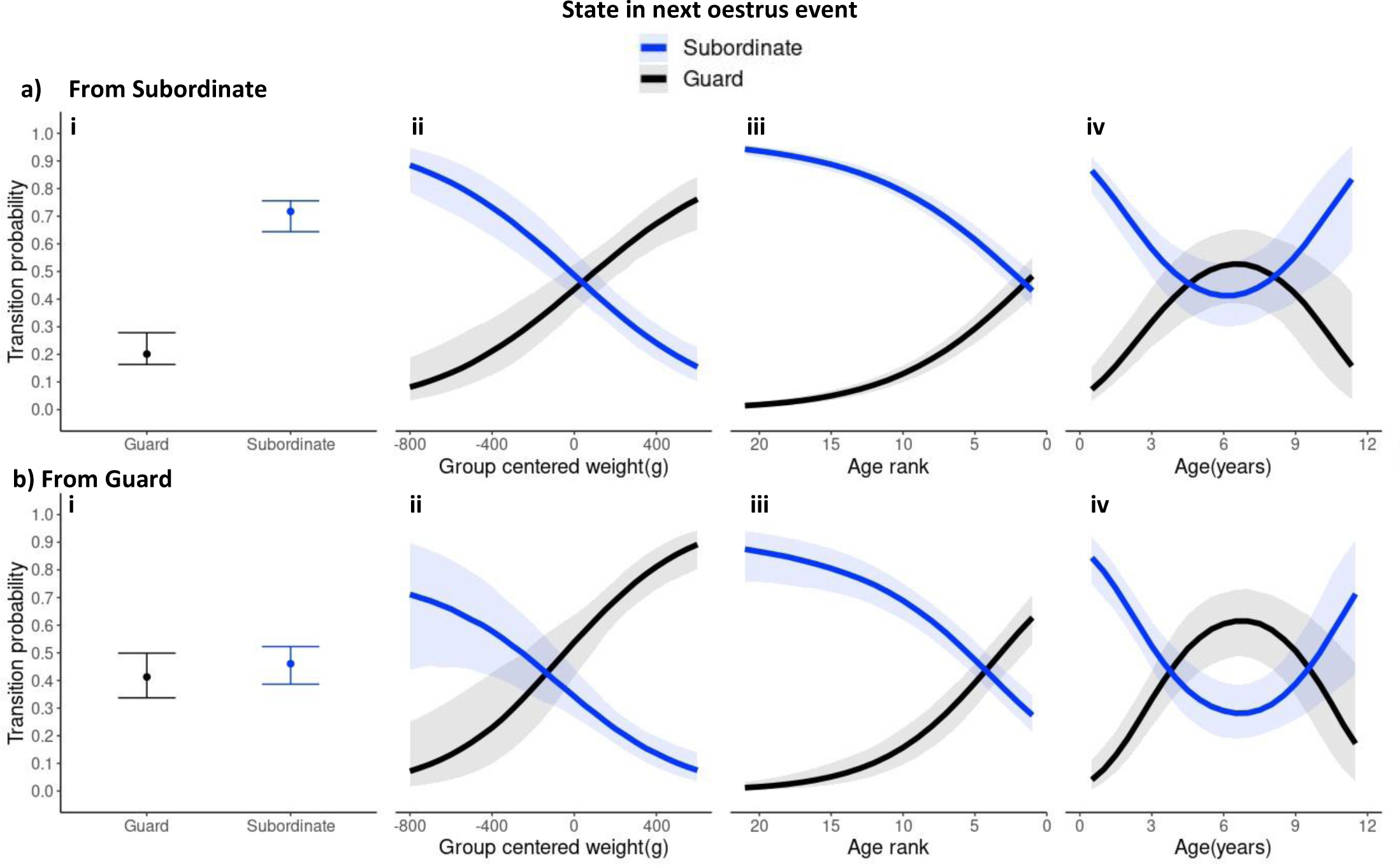
The effect of group centred weight (ii), age rank (iii), and age (iv) on the probability of acting as a subordinate or guard in the next oestrus event, given they acted as a subordinate (a) or guard (b) in the previous oestrus event. Lines represent the mean posterior probability and ribbons are credible intervals. Black lines and ribbons correspond to transitions to an active guarding role, and blue transitions to an inactive subordinate role. Panel i) shows mean (points) and credible intervals (standard error bars) for all 4 transitions for reference based on a null model with no covariates.

Overall age rank and group centered weight had similar effects on transition probabilities (Figure 1ii,iii). There was a significant linear effect whereby as young mongooses moved up in weight and age rank in the group they became more likely to gain a guarding position and less likely to stay in a subordinate position (supplementary table S1, model 1a). Parity between the probabilities of staying as a subordinate or gaining a guarding role was reached at the oldest age ranks (1 and 2) and at around 250g above the group average. At young age ranks and low weights, males rarely became a guard, for example, subordinates at age rank 11, or 200g below the mean weight of other males in the group, were predicted to gain a guarding role only ∼10% of the time (mean=0.11, hci=0.14, lci=0.08 and mean=0.09, hci=0.13, lci=0.07) tending towards 0 at younger age ranks and lighter weights. Predictions of age rank and weight diverge at cases of extreme weight, whereby age rank 1 subordinates were predicted to gain a guarding role around 50% of the time (mean=0.48, hci=0.55, lci=0.41), while those of extreme weights (97.5% quantile +320.4g, max=+494.48g) had a higher predicted probability to become a guard, approaching two times out of three (weight=+450g, mean=0.67, hci=0.77, lci=0.58).

Similarly, there was a significant linear effect of weight and age rank for keeping or losing a guarding role; heavier and older age rank males were more likely to keep a guarding position once obtained, and less likely to become an inactive subordinate (supplementary table S1, model 1a). Parity for keeping as opposed to losing a guarding role was reached at age rank 4 and +125g, younger and lighter than when parity was reached for staying or moving out from a subordinate role (age rank 1 or 2 and +250g). At the low end of probabilities males had ∼10% probability of keeping their guarding position at age rank 12 and weight-250g (mean=0.1, hci=0.16, lci=0.05 and mean= 0.1, hci=0.17, lci=0.05). Guards of age rank older than 4 were more likely to hold than lose their guarding role with age rank 1 males, who as a subordinate would have had parity in their transition probabilities, having a 63% probability of keeping their guarding role (lci=0.53, hci=0.75). Guards of extreme weights (97.5% quantile = +382.59g, max=+520.63g) had a predicted probability of keeping a guarding role exceeding 80% (weight=+450, mean= 0.81, hci=0.89, lci=0.7), higher than predicted at the highest age ranks.

Age had a significant quadratic effect on all transition probabilities (supplementary table S1, model 1b). The probability of gaining and keeping a guarding role mirrored the effect of age rank (Figure 1aiii, biii), with males tending to remain reproductively inactive well past sexual maturity (1 year) until reaching a peak of guarding at 6.5 years of age (subordinate to guard: mean=0.57, hci=0.67, lci=0.48 and staying guard: mean=0.65, hci=0.75, lci=0.56), after which probabilities of guarding decreased continuously as males aged. The switch point for the quadratic effect at 6.5 years was just below the mean age of age rank 1 males in oestrus events (mean=7.25, IQR-=3.25), suggesting an ageing effect was masked in age rank 1 males by groups that lack older males. As the probability of gaining or holding a guarding position trended down with age, parity between transitions to a subordinate and guarding role was reached at around 8.5 years (from subordinate) and 9.5 years (from guarding) (figure 1iv). At extreme ages (subordinate: 97.5% quantile = 10.24, maximum = 11.8, guards: 97.5% quantile = 10.13 years, maximum = 11.36 years) transition probabilities to gain or keep a guarding role were as low as 20% and 30% respectively (subordinate to guard: age=10.5 years, mean = 0.19, hci=0.37, lci=0.09, stay guard: age =10.5 years, mean =0.3, hci=0.53, lci=0.13). Beyond 8 years of age, increasing age corresponded to a decrease in weight of males (Figure S1) which may correspond with the decreased probability to gain and keep a guarding role at extreme ages and low group centered weights. A linear decline past the quadratic switch points was further verified by running separate models on a truncated dataset (See supplementary material).

The probability to maintain or transition into guarding roles reflected the probability of siring offspring. Older age rank and heavier males were significantly more likely to sire a given pup (supplementary table S1, model 2a), while age had a significant negative quadratic effect (supplementary table S1, 2b) with a decline in the probability of siring at old ages, past 7 years old (see supplementary for more detail, figure S2d).

The mean observed weight for a male mongoose was 1455.99 ± 4.64 s.e.m. Guards lost on average 2.36% (Figure S3; hci=-1.46%, lci=-3.33%) of their body weight 2.63% (∼40g), significantly more than subordinates (supplementary table S1, model 3) for which net change was not significantly different from 0 (Figure S3; mean =+0.21%, hci = +0.92%, lci =-0.53%-posterior effect overlaps 0).

Mean survival between oestrus events was not significantly different between guards (mean=0.075, hci=0.14, lci=0.40) and subordinates (mean=0.71, hci=0.131, lci=0.039) (Figure 4a). There was a significant negative effect of weight on mortality probabilities for guards but not subordinates (supplementary table S1, model 4a,4b). Below the average weight of males, guards started to have a higher mortality probability than subordinates (Figure 3b), although only at particularly low weights was this difference statistically significant (−450g guard: mean = 0.3, hci=0.60, lci=0.10;-450g subordinate: mean= 0.05, hci=0.10, lci=0.02). The mortality cost in low weight guards was reflected in the raw data; relative to the weight of other males in the group, 29/200 (14.5%) of guards of below average weight, and 2/908 (8%) of average and above weight, died before the next oestrus event. In contrast, 80/1190 (6.7%) of subordinates of below average weight, similar to 80/976 (8.2%) of average and above weight, died before the next oestrus event.

Mortality increased as mongooses moved into older age ranks (supplementary table S1). When simulating mortality probabilities, the effects of each state overlapped at all age rank values (Figure 2c). When simulated overall, increases in morality with age rank were small with a <5% increase in mortality from the youngest to the oldest age ranks for subordinates and negligible increase for guards (Figure 2c). There was no significant quadratic effect of age on mortality, and the linear effect of age mirrored age rank (supplementary table S1, Figure 2d).

**Figure 2:**
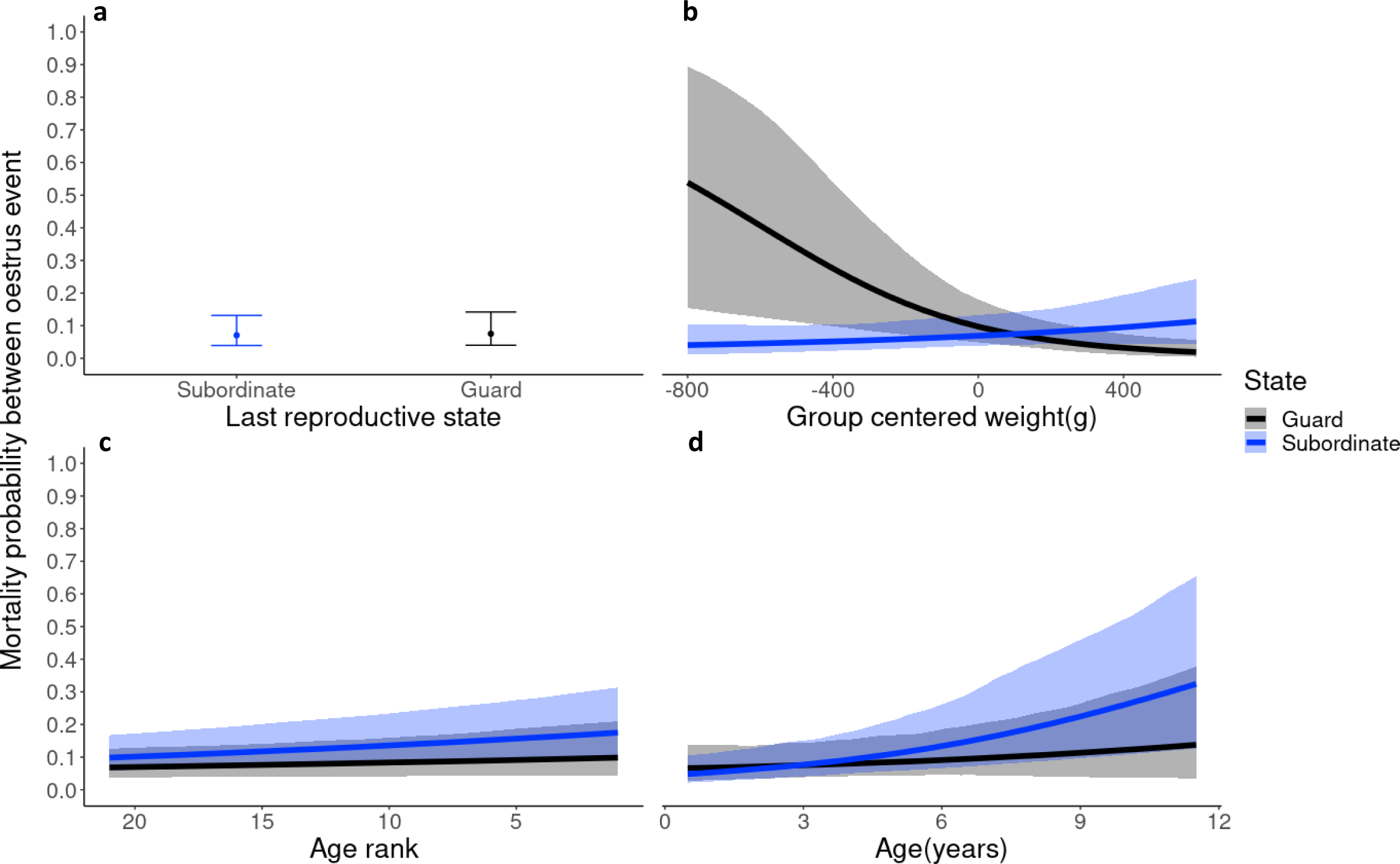
The predicted effect of male weight (b), age rank (c), and age (d) on between-oestrus event mortality probability for guards and subordinates. esent the mean posterior probability and ribbons are credible intervals. Panel a shows mean (points) and credible intervals (standard error all subordinates and guards for reference based on a null model with no covariates.

## Discussion

We found that males largely delayed reproductive activity until they became older and heavier relative to rival groups members. The oldest and the heaviest males in groups were the most reproductively active in guarding roles. Males that reached old ages (which is associated with weight declines) became less reproductively active, suggesting an ageing effect. Paternity share mirrored reproductive activity in the group, with the heaviest and oldest males in groups dominating paternity share of group litters. Reproductive activity did impose significant costs on male banded mongooses. Reproductively active guards lost weight compared to inactive subordinates who on average had no change in weight over the oestrus period. Mortality costs of reproductive activity were the greatest for lower weight males in the group, those in the poorest condition and at a RHP disadvantage to rivals.

Our finding of weight loss costs aligns with those previously reported [54], such as in reindeer (*Rangier tarandus*) during rutts [72], and other cases of mate-guarding such as in *Sceloporus virgatus* lizards [73]. The weight loss observed in our study is likely a result of the energetic demands of actively guarding females and the time spent away from foraging activities, as suggested by other studies [54,74]. Mortality costs of reproductive activity also increased as the relative weight compared to other males in the group decreased, yet only at below average weights were guard predicted to die more often than subordinates between oestrus events. Higher mortality at low weights indicates condition-dependent reproductive costs [9,10]. Condition-dependent costs align with past evidence in other systems of higher mortality costs [6,13,75] or long-term fecundity costs [15,57] of reproductive activity in lower condition breeders. Further, higher mortality costs as the weight disadavantage compared to rivals increases indicates that the social costs of conlict were raised as the RHP of competitors became more mismatched. The low-weight mortality cost of reproductive activity in male banded mongooses mirrors the rarity with which males gain or maintain active guarding roles at low weight in our transitional analysis, suggesting avoiding the costs of reproductive conflict shapes the reproductive life-histories of males in banded mongoose groups.

Condition-dependent costs suggest that individuals may not face the same constraints for life-history trade-offs when they are in sufficiently good condition [7,76]. Instead of trade-offs, positive relationships between reproductive demands and survival can be observed as individuals that typically invest more into reproduction are in higher condition[77]. For example, survival in *Iberolacerta cyreni* lizards had a positive rather than a negative association with high reproductive activity [78]. Similarly, longevity was associated with ovarian function in *Bombus teretris* honeybees [79]. Without experimental manipulation or control of condition, variation in individual condition can often mask the costs of reproduction [80]. Survival costs may have only been detectable in our system due to the longevity of our study, allowing relatively rare cases of constrained low-weight males engaging in reproductive activity to occur. Overall, reproductive costs in banded mongooses appear to follow the ‘big house big car’ analogy [9,10,77] where young lighter males cannot afford the costs of reproductive activity compared to older, heavier males who have accumlated enough wealth in terms of condition to afford the investment.

Exceptionally heavy males were the most reproductively active. The large condition advantage should be beneficial when considering the long-term endurance of consecutive bouts of reproductive activity. Banded mongooses reproduce throughout the year with new oestrus events often occuring 2 months after the last, soon after pups have emerged from the den [81], or more frequently where litters are unsuccesful. Frequent reproductive bouts mean that males must strive to recover rapidly from the costs of previous reproductive activity, which may lead to exhaustion going into the next reproductive bout if they have not replinished their condition in the interim period. Lower condition males should suffer more from reproductive exhaustion due to having fewer resources to recover from previous reproductive activity. For example, experimental manipulations of the mating history of mosquito-fish (*Gambusia affinis*) revealed ageing males are unable to maintain sperm count and velocity compared to younger males during repeated bouts of reproduction [82]. Additionally, as the weight advantage a male has over rivals increases, they may better maintain a RHP advantage into future oestrus events to more easily secure guarding positions in the long term despite repeated episodes of weight loss.

Our investigation revealed adjustments in the reproductive activity of male banded mongooses consistent with reproductive queues. Males younger than the oldest age ranks or lighter than the heaviest males delay reproductive activity despite being of a mature age, preferring to stay in inactive subordinate roles. Social costs of competiting with heavy males may expain why younger and lighter males avoid reproductive activity. The high RHP individuals that dominate reproductive roles in social groups often enforce rigid social hierarchies through regular threat displays or violence [26,58,83,84]. Yet, in male banded mongooses, outside of guarding females during short oestrus events, males are rarely observed threatening or attacking other male group members, suggesting a permanent hierarchy does not need to be enforced by dominant males. When low RHP males do engage in reproductive activity they may suffer larger social costs due to a competitive disadvantage in fights, which may have contributed to the higher mortality costs of reproductive activity at low weight relative to other group members. Self-restraint when at a competitive disadvantage likely serves to avoid social costs of reproductive conflict. These males may be passively coerced into self-restraint consistant with previously identified ‘hidden threats’, where escalation to direct suppression or violent punishment by dominants is rarely seen due to rare subordinate transgression [85]. For example, experimental manipulation revealed subordinate female banded mongooses that breed asynchronously with dominants are punished by infanticide; rarely seen under natural circumstances because females almost always breed synchronously [85]. We suggest low RHP males may similarly abstain from reproductive activity to avoid aggression by dominants. Dominant males may gain from ‘hidden threats’ as it reduces the need to actively suppress subordinate reproduction which can be costly for dominants, for example females that carried out evictions in banded mongoose groups reduced their future fecundity [86]. Instead of inccuring potentially large social costs, low RHP males may typically choose to stay reproductively inactive. Inactivity allows these males to continue gaining inclusive fitness by helping relatives [87–89].

Furthermore, reproductive inactivity may avoid delaying the growth of males in order to effectively fight in the future and ultimately secure reproductive success. For example, the energetic demands of early roaming in male African striped-mice (*Rhabdomys pumilio*) reduced life-time reproductive success by delaying the age at which males acquired their own harems [56,57]. Similarly, the weight lost by guarding early in banded mongoose groups may delay the weight gain necessary to more successfully compete for guarding roles in future oestrus events, suggesting early reproductive activity could have latent fecundity costs together with the short-term survival costs we have found. Therefore, as well as avoiding social costs of reproductive conflict in the short-term, the decision of young growing males to stay inactive in queues may be reinforced by the long-term need to gain weight to successfully challenge in future oestrus events. Self-restraint by low RHP males we evidence here suggest the costs of reproductive conflict shape male reproductive life-histories.

As condition declined with age in senescing males, the probability they maintained reproductive activity, or sired offspring, similarly declined. As their RHP declines these males may become vulnerable to displacement by rivals, aligning with senescence associated exiting of reproductive roles in other animal societies. For example, Seba’s short-tailed bats (*Carollia perspicillata)* show a decline in the abiility to maintain harems as males pass 5 years of age [68]. Also, senescing males’ reluctance to engage in reproductive activity may persist due to the threat of social conflict with higher RHP rivals. However, when so close to the end of life, abstaining from costly reproductive conflict may not contribute significantly to their lifetime reproductive success in the future. In fact, according to terminal investment theory, senescing individuals should invest maximally in reproduction when it becomes clear that there are few remaining reproductive opportunities [90]. Additionally, many intermediate weight males relative to rivals remain inactive despite no significant difference in mortality costs between activity and inactivity found, which may appear suboptimal for their lifetime reproductive fitness. These seemingly cautious approaches to reproductive activity may only make sense when considering kin selected costs. Rivals are fellow productive group members and often close kin[91], so costs imposed on them due to reproductive conflict may reduce inclusive fitness.

The costs incurred through internal conflict may weaken the group as a whole. Weight loss and injuries due to conflict may mean members are not able to effectively contribute to cooperative behaviours such as offspring care, protection against predators, or fighting during intergroup wars [33]. Reduced contributions of individuals may reduce the survival and reproductive success of all group members. As such, models of the evolution of cooperative breeders centre around how conflict has been minimised [19]. Two alternative evolutionary pathways may minimise conflict and are characterised by mechanisms that are difficult to disentangle: suppression and voluntary restraint.

One pathway we have discussed is the suppression of conflict by dominants through imposing social costs on subordinates. For example in social insects, queens police reproduction in workers by promoting aggression and the consumption of the eggs of any females that attempt to reproduce [92,93]. The second pathway is kin selection, where individuals reduce reproductive conflict in order to help relatives for inclusive fitness benefits [19]. If kin selected benefits are high voluntary restraint may be selected without the need for suppression by dominants. For example, one experimental study found policing in carpenter ants (*camponotus floridanu)* was not present in incipient colonies [94]. Worker reproduction is highly damaging to incipient colonies due to a lack of caring capacity, and policing was suggested to not be necessary in these cases as workers are under high kin selection to express voluntary reproductive restraint and help care instead of reproduce themselves [94]. Without such carefully designed experiments, suppression and voluntary restraint are difficult to tease apart. Suppression may be present but not expressed unless subordinates transgress which may only rarely be observable in natural systems. This gives the appearance of voluntary restraint, such as if subordinates use cues to avoid conflict, for example pheromones or a queen’s CHC profile in social insects [92,95]. We cannot tease apart suppression versus voluntary restraint in our wild study system, but voluntary restraint could play a role in banded mongooses groups which often have overlapping generations of related males [81]. Reducing kin-selected costs of internal conflict may serve as a significant selection pressure reinforcing the prevalent reproductive inactivity observed among mature males in this study.

The reduction in internal conflict among related males in banded mongoose groups is consistent with recent ideas suggesting a key step in evolutionary transitions to higher levels of social organisation is the breaking of life history tradeoffs (i.e. major evolutionary transitions) [96]. Kin selection may favour low condition males to remain inactive in order to allow higher condition males to reproduce who face a weakened trade-off between reproduction and survival. Kin selection could also favour reduction in the social costs of internal conflict to improve the fighting ability of the group in intergroup wars, as males specifically are important in determing the outcome of conflict [61]. Of course as discussed, the reproductive inactivity of lower condition mature males we have found may simply be a result of competitive displacement from reproductive positions, or self-interested attempts by subordinates to avoid the social costs of conflict in the group, which does not require kin selection to play a role. In our wild population males are typically closely related to rival same-sex group members [81]. Without many social groups with non-related rivals as a comparison to related-rivals it will be difficult to disentangle how self interest and kin selection mould the reproductive life-histories of male banded mongooses. Experimentally manipulated group demographies are not always possible when monitoring long-term projects, as in our case, but where they are possible comparisons of reproductive activity trajectories couldallow us to understand the role kin selection plays moulding the social life-histories of males in social groups.

## Conclusion

The life-histories of male banded mongooses are shaped by the benefits of reducing costly reproductive conflict. Lower condition males remain inactive which likely serves to avoid suffering higher social costs of reproductive activity, supported by higher mortality costs found for lower weight males. These costs fit well with the queueing dynamics we have found as young, lighter males delay reproductive activity, which may allow more succesful attempts to compete with rivals in the future. As males grow, the social costs of reproductive conflict decline, and increased condition makes the energetic demands of guarding easier to bear. Reduction in the costs of conflict is reflected in more consistant engagement in reproductive activity at older, but not senescent, age classes. Kin selected benefits may play a role reducing reproductive conflict within banded mongoose groups. Disentangling how self interest and kin selection mould the reproductive life-histories of males in social groups requires future research.

## Acknowledgements

Special thanks to Nicolas Fasel for help getting started using State transition models in JAGS. Additionally thanks to current and past members of our research group for helpful discussion and input.

## References

1. Monaghan P, Metcalfe NB, Torres R. 2009 Oxidative stress as a mediator of life history trade-offs: Mechanisms, measurements and interpretation. Ecol. Lett. 12, 75–92. (doi:10.1111/j.1461-0248.2008.01258.x)

2. Speakman JR, Garratt M. 2014 Oxidative stress as a cost of reproduction: Beyond the simplistic trade-off model. BioEssays 36, 93–106. (doi:10.1002/bies.201300108)

3. Zhang Y, Hood WR. 2016 Current versus future reproduction and longevity: A re-evaluation of predictions and mechanisms. J. Exp. Biol. 219, 3177–3189. (doi:10.1242/jeb.132183)

4. Harshman LG, Zera AJ. 2007 The cost of reproduction: the devil in the details. Trends Ecol. Evol. 22, 80–86. (doi:10.1016/j.tree.2006.10.008)

5. Cody ML. 1966 A General Theory of Clutch Size. Evolution (N. Y). 20, 174–184. (doi:10.2307/2406571)

6. Kirkwood TBL, Rose MR. 1991 Evolution of senescence: late survival sacrificed for reproduction. *Philos. Trans.-R. Soc. London*, B 332, 15–24. (doi:10.1098/rstb.1991.0028)

7. Stearns SC. 1992 The Evolution of Life-histories. Oxford: Oxford University Press. (doi:10.2307/5403)

8. Pianka ER, Parker WS. 1975 Age-Specific Reproductive Tactics. Am. Nat. 109, 453–464. (doi:10.1086/283013)

9. Reznick D, Nunney L, Tessier A. 2000 Big houses, big cars, superfleas and the costs of reproduction. Trends Ecol. Evol. 15, 421–425. (doi:10.1016/S0169-5347(00)01941-8)

10. Metcalfe NB, Monaghan P. 2013 Does reproduction cause oxidative stress? An open question. Trends Ecol. Evol. 28, 347–350. (doi:10.1016/j.tree.2013.01.015)

11. Nichols HJ, Amos W, Cant MA, Bell MBV, Hodge SJ. 2010 Top males gain high reproductive success by guarding more successful females in a cooperatively breeding mongoose. Anim. Behav. 80, 649–657. (doi:10.1016/j.anbehav.2010.06.025)

12. Groenewoud F, Clutton-Brock T. 2021 Meerkat helpers buffer the detrimental effects of adverse environmental conditions on fecundity, growth and survival. J. Anim. Ecol. 90, 641–652. (doi:10.1111/1365-2656.13396)

13. Lemaître JF, Berger V, Bonenfant C, Douhard M, Gamelon M, Plard F, Gaillard JM. 2015 Early-late life trade-offs and the evolution of ageing in the wild. Proc. R. Soc. B Biol. Sci. 282. (doi:10.1098/rspb.2015.0209)

14. Lemaître JF, Gaillard JM, Pemberton JM, Clutton-Brock TH, Nussey DH. 2014 Early life expenditure in sexual competition is associated with increased reproductive senescence in male red deer. Proc. R. Soc. B Biol. Sci. 281. (doi:10.1098/rspb.2014.0792)

15. Schradin C, Lindholm AK, Johannesen J, Schoepf I, Yuen CH, König B, Pillay N. 2012 Social flexibility and social evolution in mammals: A case study of the African striped mouse (Rhabdomys pumilio). Mol. Ecol. 21, 541–553. (doi:10.1111/C65-294X.2011.05256.x)

16. Doerr ED, Doerr VAJ. 2007 Positive effects of helpers on reproductive success in the brown treecreeper and the general importance of future benefits. J. Anim. Ecol. 76, 966–976. (doi:10.1111/j.1365-2656.2007.01280.x)

17. Gilchrist JS. 2006 Reproductive success in a low skew, communal breeding mammal: The banded mongoose, Mungos mungo. Behav. Ecol. Sociobiol. 60, 854–863. (doi:10.1007/s00265-006-0229-6)

18. O’Riain MJ, Bennett NC, Brotherton PNM, McIlrath G, Clutton-Brock TH. 2000 Reproductive suppression and inbreeding avoidance in wild populations of co-operatively breeding meerkats (Suricata suricatta). Behav. Ecol. Sociobiol. 48, 471– 477. (doi:10.1007/s002650000249)

19. Taborsky M, Cant MA, Komdeur J. 2021 Conflict. In The Evolution of Social Behaviour, pp. 67–135. Cambridge University Press. (doi:10.1017/9780511894794.005)

20. Holmes MM, Goldman BD, Goldman SL, Seney ML, Forger NG. 2009 Neuroendocrinology and sexual differentiation in eusocial mammals. Front. Neuroendocrinol. 30, 519–533. (doi:10.1016/j.yfrne.2009.04.010)

21. Faulkes CG, Abbott DH, Jarvis JUM. 1991 Social suppression of reproduction in male naked mole-rats, Heterocephalus glaber. J. Reprod. Fertil. 91, 593–604. (doi:10.1530/jrf.0.0910593)

22. Zhou S, Holmes MM, Forger NG, Goldman BD, Lovern MB, Caraty A, Kalló I, Faulkes CG, Coen CW. 2013 Socially regulated reproductive development: Analysis of GnRH-1 and kisspeptin neuronal systems in cooperatively breeding naked mole-rats (Heterocephalus glaber). J. Comp. Neurol. 521, 3003–3029. (doi:10.1002/cne.23327)

23. Swift-Gallant A, Mo K, Peragine DE, Ashley Monks D, Holmes MM. 2015 Removal of reproductive suppression reveals latent sex differences in brain steroid hormone receptors in naked mole-rats, Heterocephalus glaber. Biol. Sex Differ. 6, 1–9. (doi:10.1186/s13293-015-0050-x)

24. Carlson AA, Young AJ, Russell AF, Bennett NC, McNeilly AS, Clutton-Brock T. 2004 Hormonal correlates of dominance in meerkats (Suricata suricatta). Horm. Behav. 46, 141–150. (doi:10.1016/j.yhbeh.2004.01.009)

25. Montgomery TM, Pendleton EL, Smith JE. 2018 Physiological mechanisms mediating patterns of reproductive suppression and alloparental care in cooperatively breeding carnivores. Physiol. Behav. 193, 167–178. (doi:10.1016/j.physbeh.2017.11.006)

26. Thompson FJ, Donaldson L, Johnstone RA, Field J, Cant MA. 2014 Dominant aggression as a deterrent signal in paper wasps. Behav. Ecol. 25, 706–715. (doi:10.1093/beheco/aru063)

27. Smith JM, Harper D. 2003 Animal signals. Oxford University Press.

28. Ross CT, Hooper PL, Smith JE, Jaeggi A V, Alden E, Gavrilets S. 2023 Reproductive inequality in humans and other mammals. 120, 1–12. (doi:10.1073/pnas.2220124120/-/DCSupplemental.Published)

29. Fitzgerald LM et al. 2022 Rank change and growth within social hierarchies of the orange clownfish, Amphiprion percula. Mar. Biol. 169, 128. (doi:10.1007/s00227-022-04117-9)

30. Stephens PA, Russell AF, Young AJ, Sutherland WJ, Clutton-Brock TH. 2005 Dispersal, eviction, and conflict in meerkats (Suricata suricatta): An evolutionarily stable strategy model. Am. Nat. 165, 120–35. (doi:10.1086/426597)

31. Cant MA, Hodge SJ, Bell MBV, Gilchrist JS, Nichols HJ. 2010 Reproductive control via eviction (but not the threat of eviction) in banded mongooses. Proc. R. Soc. B Biol. Sci. 277. (doi:10.1098/rspb.2009.2097)

32. Iwasa Y, Yamaguchi S. 2022 On the role of eviction in group living sex changers. Behav. Ecol. Sociobiol. 76, 49. (doi:10.1007/s00265-022-03159-9)

33. Reeve HK, Hölldobler B. 2007 The emergence of a superorganism through intergroup competition. Proc. Natl. Acad. Sci. U. S. A. 104, 9736–9740. (doi:10.1073/pnas.0703466104)

34. Cant MA, English S. 2006 Stable group size in cooperative breeders: The role of inheritance and reproductive skew. Behav. Ecol. 17, 560–568. (doi:10.1093/beheco/arj065)

35. Bridge C, Field J. 2007 Queuing for dominance: Gerontocracy and queue-jumping in the hover wasp Liostenogaster flavolineata. Behav. Ecol. Sociobiol. 61, 1253–1259. (doi:10.1007/s00265-007-0355-9)

36. Bergmüller R, Taborsky M. 2005 Experimental manipulation of helping in a cooperative breeder: Helpers ‘pay to stay’ by pre-emptive appeasement. Anim. Behav. 69, 19–28. (doi:10.1016/j.anbehav.2004.05.009)

37. Reddon AR, Ruberto T, Reader SM. 2021 Submission signals in animal groups. Behaviour 159, 1–20. (doi:10.1163/1568539X-bja10125)

38. Wong MYL, Buston PM, Munday PL, Jones GP. 2007 The threat of punishment enforces peaceful cooperation and stabilizes queues in a coral-reef fish. Proc. R. Soc. B Biol. Sci. 274. (doi:10.1098/rspb.2006.0284)

39. Hamilton IM, Heg D. 2008 Sex differences in the effect of social status on the growth of subordinates in a co-operatively breeding cichlid. J. Fish Biol. 72, 1079–1088. (doi:10.1111/j.1095-8649.2007.01787.x)

40. Mitchell J, Kyabulima S, Businge R, Cant MA, Nichols HJ. 2018 Kin discrimination via odour in the cooperatively breeding banded mongoose. R. Soc. Open Sci. 5. (doi:10.1098/rsos.171798)

41. Schneider TC, Kappeler PM. 2014 Social systems and life-history characteristics of mongooses. Biol. Rev. 89, 173–198. (doi:10.1111/brv.12050)

42. Dyble M, Clutton-Brock TH. 2020 Contrasts in kinship structure in mammalian societies. Behav. Ecol. 31, 971–977. (doi:10.1093/BEHECO/ARAA043)

43. Hamilton WD. 1964 The genetical evolution of social behavior. I and II. J. Theor. Biol. 7, 1–16.

44. Downing PA, Cornwallis CK, Griffin AS. 2015 Sex, long life and the evolutionary transition to cooperative breeding in birds. Proc. R. Soc. B Biol. Sci. 282. (doi:10.1098/rspb.2015.1663)

45. Kreider JJ, Kramer BH, Komdeur J, Pen I. 2022 The evolution of ageing in cooperative breeders. Evol. Lett. 6, 450–459. (doi:10.1002/evl3.307)

46. Nielsen MLK et al. 2021 A long postreproductive life span is a shared trait among genetically distinct killer whale populations. Ecol. Evol. 11, 9123–9136. (doi:10.1002/ece3.7756)

47. Johnstone RA, Cant MA. 2010 The evolution of menopause in cetaceans and humans: The role of demography. Proc. R. Soc. B Biol. Sci. 277. (doi:10.1098/rspb.2010.0988)

48. Thompson FJ, Marshall HH, Sanderson JL, Vitikainen EIK, Nichols HJ, Gilchrist JS, Young AJ, Hodge SJ, Cant MA. 2016 Reproductive competition triggers mass eviction in cooperative banded mongooses. Proc. R. Soc. B Biol. Sci. 283. (doi:10.1098/rspb.2015.2607)

49. Ellis S et al. 2022 Patterns and consequences of age-linked change in local relatedness in animal societies. *Nat*. Ecol. Evol. 6, 1766–1776. (doi:10.1038/s41559-022-01872-2)

50. Trillmich F, Geißler E, Guenther A. 2019 Senescence and costs of reproduction in the life history of a small precocial species. Ecol. Evol., 7069–7079. (doi:10.1002/ece3.5272)

51. Croft DP et al. 2017 Reproductive Conflict and the Evolution of Menopause in Killer Whales. Curr. Biol. 27, 298–304. (doi:10.1016/j.cub.2016.12.015)

52. Lahdenperä M, Gillespie DOS, Lummaa V, Russell AF. 2012 Severe intergenerational reproductive conflict and the evolution of menopause. Ecol. Lett. 15, 1283–1290. (doi:10.1111/j.1461-0248.2012.01851.x)

53. Friesen CR, De Graaf SP, Olsson M. 2019 The relationship of body condition, superoxide dismutase, and superoxide with sperm performance. Behav. Ecol. 30, 1351–1363. (doi:10.1093/beheco/arz086)

54. Ord TJ. 2021 Costs of territoriality: a review of hypotheses, meta-analysis, and field study. Oecologia 197, 615–631. (doi:10.1007/s00442-021-05068-6)

55. Sharick JT, Vazquez-Medina JP, Ortiz RM, Crocker DE. 2015 Oxidative stress is a potential cost of breeding in male and female northern elephant seals. Funct. Ecol. 29, 367–376. (doi:10.1111/1365-2435.12330)

56. Rimbach R, Blanc S, Zahariev A, Pillay N, Schradin C. 2019 Daily energy expenditure of males following alternative reproductive tactics: Solitary roamers spend more energy than group-living males. Physiol. Behav. 199, 359–365. (doi:10.1016/j.physbeh.2018.12.003)

57. Kanyile SN, Pillay N, Schradin C. 2021 Bachelor groups form due to individual choices or environmental disrupters in African striped mice. Anim. Behav. 182, 135–143. (doi:10.1016/j.anbehav.2021.10.005)

58. Cant MA. 2000 Social control of reproduction in banded mongooses. Anim. Behav. 59, 147–158. (doi:10.1006/anbe.1999.1279)

59. Cant MA, Vitikainen E, Nichols HJ. 2013 Demography and Social Evolution of Banded Mongooses. Adv. Study Behav. 45, 407–445. (doi:10.1016/B978-0-12-407186-5.00006-9)

60. Vitikainen EIK, Thompson FJ, Marshall HH, Cant MA, Cant MA. 2019 Live long and prosper: durable benefits of early-life care in banded mongooses. 374.

61. Green PA, Thompson FJ, Cant MA. 2022 Fighting force and experience combine to determine contest success in a warlike mammal. Proc. Natl. Acad. Sci. 119, e2119176119. (doi:10.1073/pnas.2119176119)

62. Fasel NJ, Wesseling C, Fernandez AA, Vallat A, Glauser G, Helfenstein F, Richner H. 2017 Alternative reproductive tactics, sperm mobility and oxidative stress in Carollia perspicillata (Seba’s short-tailed bat). Behav. Ecol. Sociobiol. 71, 11. (doi:10.1007/s00265-016-2251-7)

63. Campos FA, Villavicencio F, Archie EA, Colchero F, Alberts SC. 2020 Social bonds, social status and survival in wild baboons: a tale of two sexes. Philos. Trans. R. Soc. B Biol. Sci. 375. (doi:10.1098/rstb.2019.0621)

64. Nakagawa S. 2015 Missing data: mechanisms, methods, and messages. Ecol. Stat. Contemp. Theory Appl., 81–105.

65. Gilchrist JS. 2004 Pup escorting in the communal breeding banded mongoose: Behavior, benefits, and maintenance. Behav. Ecol. 15, 952–960. (doi:10.1093/beheco/arh071)

66. Kéry M, Schaub M, Beissinger SR. 2012 Bayesian Population Analysis using WinBUGS A hierarchical perspective. See http://www.elsevierdirect.com.

67. Marin X. 2021 ggmcmc: Tools for Analyzing MCMC Simulations from Bayesian Inference. See https://cran.r-project.org/package=ggmcmc.

68. Fasel N, Saladin V, Richner H. 2016 Alternative reproductive tactics and reproductive success in male Carollia perspicillata (Seba’s short-tailed bat). J. Evol. Biol. 29, 2242– 2255. (doi:10.1111/jeb.12949)

69. Freckleton RP. 2011 Dealing with collinearity in behavioural and ecological data: Model averaging and the problems of measurement error. Behav. Ecol. Sociobiol. 65, 91–101. (doi:10.1007/s00265-010-1045-6)

70. Youngflesh C, Che-Castaldo C, Hardy T. In press. MCMCvis: Tools to Visualize, Manipulate, and Summarize MCMC Output. See https://cran.r-project.org/package=MCMCvis.

71. Kellner K, Meredith M. 2024 jagsUI: A Wrapper Around ‘rjags’ to Streamline ‘JAGS’ Analyses. See https://cran.r-project.org/package=jagsUI.

72. Tennenhouse EM, Weladji RB, Holand Ø, Røed KH, Nieminen M. 2011 Mating group composition influences somatic costs and activity in rutting dominant male reindeer (Rangifer tarandus). Behav. Ecol. Sociobiol. 65, 287–295. (doi:10.1007/s00265-010-1043-8)

73. Abell AJ. 2000 Costs of reproduction in male lizards, Sceloporus virgatus. Oikos 88, 630–640. (doi:10.1034/j.1600-0706.2000.880320.x)

74. Ancona S, Drummond H, Zaldívar-Rae J. 2010 Male whiptail lizards adjust energetically costly mate guarding to male-male competition and female reproductive value. Anim. Behav. 79, 75–82. (doi:10.1016/j.anbehav.2009.10.005)

75. Ritchot Y, Festa-Bianchet M, Coltman D, Pelletier F. 2021 Determinants and long-term costs of early reproduction in males of a long-lived polygynous mammal. Ecol. Evol. 11, 6829–6845. (doi:10.1002/ece3.7530)

76. Williams TD. 2018 Physiology, activity and costs of parental care in birds. J. Exp. Biol. 221, eb169433. (doi:10.1242/jeb.169433)

77. Zera AJ, Harshman LG. 2001 The physiology of life history trade-offs in animals. Annu. Rev. Ecol. Syst. 32, 95–126. (doi:10.1146/annurev.ecolsys.32.081501.114006)

78. Salvador A, Díaz JA, Veiga JP, Bloor P, Brown RP. 2008 Correlates of reproductive success in male lizards of the alpine species Iberolacerta cyreni. Behav. Ecol. 19, 169–176. (doi:10.1093/beheco/arm118)

79. Blacher P, Huggins TJ, Bourke AFG. 2017 Evolution of ageing, costs of reproduction and the fecundity–longevity trade-off in eusocial insects. Proc. R. Soc. B Biol. Sci. 284. (doi:10.1098/rspb.2017.0380)

80. Cox RM, Wittman TN, Calsbeek R. 2022 Reproductive trade-offs and phenotypic selection change with body condition, but not with predation regime, across island lizard populations. J. Evol. Biol. 35, 365–378. (doi:10.1111/jeb.13926)

81. Cant MA, Nichols HJ, Thompson FJ, Vitikainen E. 2016 Banded mongooses: Demography, life history, and social behavior. In *Cooperative Breeding in Vertebrates: Studies of Ecology*, Evolution, and Behavior, pp. 318–337. (doi:10.1017/CBO9781107338357.019)

82. Aich U, Head ML, Fox RJ, Jennions MD. 2021 Male age alone predicts paternity success under sperm competition when effects of age and past mating effort are experimentally separated. Proc. R. Soc. B Biol. Sci. 288, 1955. (doi:10.1098/rspb.2021.0979)

83. Fichtel C, Kraus C, Ganswindt A, Heistermann M. 2007 Influence of reproductive season and rank on fecal glucocorticoid levels in free-ranging male Verreaux’s sifakas (Propithecus verreauxi). Horm. Behav. 51, 640–648. (doi:10.1016/j.yhbeh.2007.03.005)

84. Young AJ, Carlson AA, Monfort SL, Russell AF, Bennett NC, Clutton-Brock T. 2006 Stress and the suppression of subordinate reproduction in cooperatively breeding meerkats. Proc. Natl. Acad. Sci. U. S. A. 103, 12005–12010. (doi:10.1073/pnas.0510038103)

85. Cant MA, Nichols HJ, Johnstone RA, Hodge SJ. 2013 Policing of reproduction by hidden threats in a cooperative mammal. PNAS 111, 326–330. (doi:10.1073/pnas.1312626111/-/DCSupplemental.www.pnas.org/cgi/doi/10.1073/pnas.1312626111)

86. Bell MBV, Nichols HJ, Gilchrist JS, Cant MA, Hodge SJ. 2012 The cost of dominance: Suppressing subordinate reproduction affects the reproductive success of dominant female banded mongooses. Proc. R. Soc. B Biol. Sci. 279, 619–624. (doi:10.1098/rspb.2011.1093)

87. West SA, Cooper GA, Ghoul MB, Griffin AS. 2021 Ten recent insights for our understanding of cooperation. *Nat*. Ecol. Evol. 5. (doi:10.1038/s41559-020-01384-x)

88. Hamilton WD. 1970 Selfish and spiteful behaviour in an evolutionary model. Nature 228, 1218–1220.

89. Frank SA. 2003 Perspective: Repression of competition and the evolution of cooperation. Evolution (N. Y). 57, 693–705. (doi:10.1111/j.0014-3820.2003.tb00283.x)

90. Duffield KR, Bowers EK, Sakaluk SK, Sadd BM. 2017 A dynamic threshold model for terminal investment. Behav. Ecol. Sociobiol. 71, 185. (doi:10.1007/s00265-017-2416-z)

91. Nichols HJ, Jordan NR, Jamie GA, Cant MA, Hoffman JI. 2012 Fine-scale spatiotemporal patterns of genetic variation reflect budding dispersal coupled with strong natal philopatry in a cooperatively breeding mammal. Mol. Ecol. 21. (doi:10.1111/mec.12015)

92. Le Conte Y, Hefetz A. 2008 Primer pheromones in social hymenoptera. Annu. Rev. Entomol. 53, 523–542. (doi:10.1146/annurev.ento.52.110405.091434)

93. Wenseleers T, Hart AG, Ratnieks FLW. 2004 When Resistance Is Useless: Policing and the Evolution of Reproductive Acquiescence in Insect Societies. Am. Nat. 164, 154–167. (doi:10.1086/425223)

94. Moore D, Liebig J. 2013 Reproductive restraint without policing in early stages of a social insect colony. Anim. Behav. 85, 1323–1328. (doi:10.1016/j.anbehav.2013.03.022)

95. Smith AA, Hölldober B, Liebig J. 2009 Cuticular Hydrocarbons Reliably Identify Cheaters and Allow Enforcement of Altruism in a Social Insect. Curr. Biol. 19, 78–81. (doi:10.1016/j.cub.2008.11.059)

96. Bourrat P, Doulcier G, Rose CJ, Rainey PB, Hammerschmidt K. 2022 Tradeoff breaking as a model of evolutionary transitions in individuality and limits of the fitness-decoupling metaphor. Elife 11, e73715. (doi:10.7554/eLife.73715)

