## Supplementary material for "Reproductive restraint to avoid the costs of reproductive conflict in a cooperatively breeding mammal"

**Supplementary material for reproductive restraint in male banded mongooses**

**Supplementary methods**

*Weight inputation*

Weight is correlated with age in banded mongooses (Figure S1), and 93.75% of males (7/112) that required imputed weights had some history of weight collection allowing imputation to track individual weight trajectories using random slopes. Missing data was inputted using a linear mixed effect model fitted with quadratic age, random intercept (group id), and random slope on male id with quadratic age using the lme4 r package. The model was run using 48500 weights collected on the 314 males of known age above 180 days old collected since 2000. This included all males that required inputted weights. The quadratic term was justified to account for increases in age as individuals grew, and decreases in weight with senescence. This was validated by model comparisons with a linear model by likelihood ratio tests (Quadratic AIC = 28807, Linear AIC = 56608, Null AIC = 86050). Missing weights were extracted from the sampling distribution predicted from the above model simulated 10,000 times using the PredictInterval function (merTools r package).

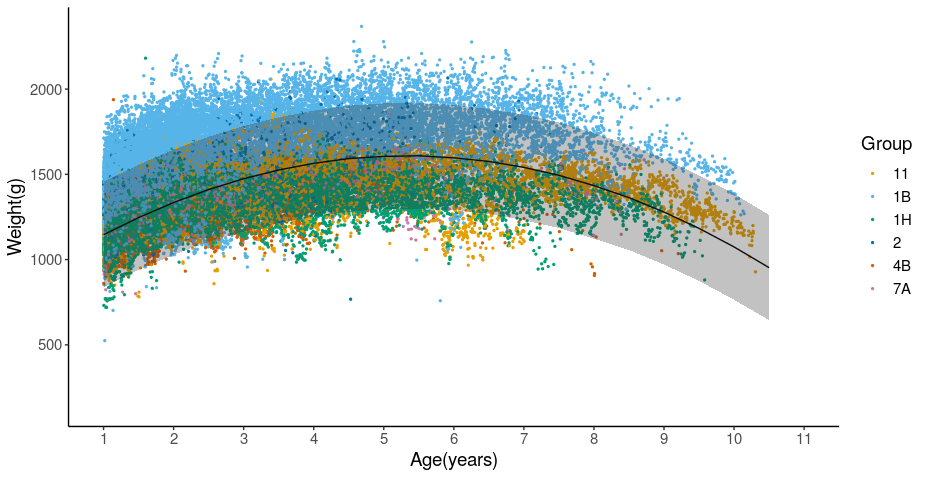

Figure S1: The relationship between age (years) and bodymass of males. Dots represent historically collected weights from 6 groups in the Mweya study population. Line and ribbon represent the output of a General Linear Model for age predicting weight with group as a random effect. Data show that weight increased with age until around 6 years of age, and then decreased.

**Supplementary results**

**Bayesian model output for all models in this study**

**Table S1**

| Summary of bayesianan general linear models fitted in this study | | | | | | | | | |
| --- | --- | --- | --- | --- | --- | --- | --- | --- | --- |
| **Model** | **Term** | **effect** | **sd** | **2.5%** | **50.0%** | **97.5%** | **Rhat** | **f** | **Overlap0** |
| 1a | Constant(StaySub) | 0.06 | 0.17 | -0.26 | 0.06 | 0.44 | 1.01 | 0.63 | Yes |
|  | Constant(SubToGuard) | 0.11 | 0.28 | -0.44 | 0.11 | 0.74 | 1 | 0.7 | Yes |
|  | Constant(GuardToSub) | -0.56 | 0.23 | -1.01 -0.58 | | -0.1 | 1 | 0.99 | - |
|  | Constant(StayGuard) | 0.81 | 0.27 | 0.29 | 0.83 | 1.28 | 1 | 1 | + |
|  | **AgeRank(StaySub)** | 0.15 | 0.02 | 0.11 | 0.15 | 0.18 | 1 | 1 | + |
|  | **AgeRank(SubToGuard)** | -0.17 | 0.02 | -0.21 | -0.17 | -0.13 | 1 | 1 | - |
|  | **AgeRank(GuardToSub)** | 0.2 | 0.03 | 0.14 | 0.2 | 0.27 | 1.01 | 1 | + |
|  | **AgeRank(StayGuard)** | -0.23 | 0.03 | -0.29 | -0.23 | -0.16 | 1 | 1 | - |
|  | **Weight(StaySub)** | -0.52 | 0.09 | -0.71 | -0.52 | -0.36 | 1 | 1 | - |
|  | **Weight(SubToGuard)** | 0.58 | 0.09 | 0.4 | 0.58 | 0.76 | 1 | 1 | + |
|  | **Weight(GuardToSub)** | -0.61 | 0.14 | -0.88 | -0.6 | -0.35 | 1 | 1 | - |
|  | **Weight(StayGuard)** | 0.73 | 0.15 | 0.45 | 0.72 | 1.02 | 1.01 | 1 | + |
|  | Random effect | Group | Male.id | Oestrus.event | |  |  |  |  |
|  | sd | 0.102 | 0.122 | 0.397 |  |  |  |  |  |
| 1b | Constant(StaySub) | 0.53 | 0.18 | 0.16 | 0.56 | 0.84 | 1.01 | 0.99 | + |
|  | Constant(SubToGuard) | -0.32 | 0.31 | -0.86 | -0.36 | 0.42 | 1 | 0.85 | Yes |
|  | Constant(GuardToSub) | 0.3 | 0.22 | -0.08 | 0.3 | 0.72 | 1.01 | 0.94 | +* |
|  | Constant(StayGuard) | -0.18 | 0.23 | -0.64 | -0.18 | 0.3 | 1.01 | 0.79 | Yes |
|  | **Age(StaySub)** | -0.94 | 0.13 | -1.2 | -0.93 | -0.69 | 1.01 | 1 | - |
|  | **Age(SubToGuard)** | 1.09 | 0.14 | 0.82 | 1.1 | 1.37 | 1 | 1 | + |
|  | **Age(GuardToSub)** | -1.25 | 0.19 | -1.63 | -1.26 | -0.86 | 1 | 1 | - |
|  | **Age(StayGuard)** | 1.3 | 0.19 | 0.92 | 1.31 | 1.71 | 1 | 1 | + |
|  | **Age^2(StaySub)** | 0.36 | 0.07 | 0.22 | 0.36 | 0.5 | 1.01 | 1 | + |
|  | **Age^2(SubToGuard)** | -0.43 | 0.07 | -0.57 | -0.43 | -0.27 | 1.01 | 1 | - |
|  | **Age^2(GuardToSub)** | 0.44 | 0.09 | 0.26 | 0.44 | 0.63 | 1 | 1 | + |
|  | **Age^2(StayGuard)** | -0.44 | 0.1 | -0.63 | -0.44 | -0.25 | 1 | 1 | - |
|  | **Weight(StaySub)** | -0.41 | 0.09 | -0.59 | -0.41 | -0.23 | 1 | 1 | - |
|  | **Weight(SubToGuard)** | 0.43 | 0.1 | 0.24 | 0.43 | 0.64 | 1.01 | 1 | + |
|  | **Weight(GuardToSub)** | -0.49 | 0.15 | -0.77 | -0.5 | -0.2 | 1 | 1 | - |
|  | **Weight(StayGuard)** | 0.6 | 0.16 | 0.29 | 0.59 | 0.91 | 1.01 | 1 | + |
|  | Random effect | Group | Male.id | Oestrus.event |  |  |  |  |  |
|  | sd | 0.129 | 0.12 | 0.407 |  |  |  |  |  |
| 2a | Constant | -1.72 | 0.19 | -2.09 | -1.72 | -1.36 | 1 | 1 | - |
|  | **Weight** | 0.81 | 0.11 | 0.58 | 0.81 | 1.02 | 1 | 1 | + |
|  | **AgeRank** | -0.34 | 0.03 | -0.41 | -0.34 | -0.28 | 1 | 1 | - |
|  | Random effect | Group | Male.id | Oestrus.event |  |  |  |  |  |
|  | sd | 0.0264 | 0.539 | 0.005 |  |  |  |  |  |
| 2b | Constant | -3.22 | 0.25 | -3.69 | -3.22 | -2.75 | 1.01 | 1 | - |
|  | **Weight** | 0.64 | 0.13 | 0.39 | 0.64 | 0.88 | 1 | 1 | + |
|  | **Age** | 1.88 | 0.17 | 1.56 | 1.87 | 2.23 | 1 | 1 | + |
|  | **Age^2** | -0.6 | -0.08 | -0.75 | -0.6 | -0.46 | 1 | 1 | - |
|  | Random effect | Group | Male.id | Oestrus.event |  |  |  |  |  |
|  | sd | 0.469 | 0.558 | 0.008 |  |  |  |  |  |
| 3 | Constant | 0.25 | 0.36 | -0.46 | 0.23 | 1.01 | 1.02 | 0.74 | Yes |
|  | **State(Guard)** | -2.63 | 0.41 | -3.51 | -2.62 | -1.86 | 1.01 | 1 | - |
|  | Random effect | Group | Male.id | Oestrus.event |  |  |  |  |  |
|  | sd | 0.45 | 0.109 | 0.359 |  |  |  |  |  |
| 4a | Constant(Sub) | -1.63 | 0.33 | -2.2 | -1.64 | -0.97 | 1.01 | 1 | - |
|  | Constant(Guard) | -1.58 | 0.41 | -2.33 | -1.6 | -0.75 | 1.01 | 1 | - |
|  | **AgeRank(Sub)** | -0.16 | 0.04 | -0.23 | -0.16 | -0.08 | 1.01 | 1 | - |
|  | **AgeRank(Guard)** | -0.15 | 0.06 | -0.29 | -0.15 | -0.04 | 1.01 | 1 | - |
|  | Weight(Sub) | -0.16 | 0.14 | -0.43 | -0.16 | 0.1 | 1 | 0.85 | Yes |
|  | **Weight(Guard)** | -0.81 | 0.23 | -1.3 | -0.79 | -0.4 | 1.01 | 1 | - |
|  | Time(Sub)* | 0.35 | 0.21 | -0.01 | 0.33 | 0.81 | 1 | 0.97 | +* |
|  | **Time(Guard)** | 0.63 | 0.25 | 0.16 | 0.61 | 1.13 | 1 | 1 | + |
|  | Random effect | Group | Male.id | Oestrus.event |  |  |  |  |  |
|  | sd | 0.105 | 0.132 | 0.953 |  |  |  |  |  |
| 4b | Constant(Sub) | -2.39 | 0.35 | -2.98 | -2.41 | -1.61 | 1 | 1 | - |
|  | Constant(Guard) | -2.17 | 0.38 | -2.9 | -2.21 | -1.42 | 1 | 1 | - |
|  | **Age(Sub)** | 0.59 | 0.23 | 0.24 | 0.55 | 1.14 | 1.02 | 1 | + |
|  | Age(Guard)* | 0.42 | 0.25 | 0 | 1 | 1.02 | 1.02 | 0.97 | +* |
|  | Weight(Sub) | -0.03 | 0.14 | -0.28 | -0.04 | 0.26 | 1 | 0.59 | Yes |
|  | **Weight(Guard)** | -0.75 | 0.23 | -1.21 | -0.74 | -0.31 | 1 | 1 | - |
|  | **Time(Sub)** | 0.51 | 0.25 | 0.09 | 0.49 | 1.01 | 1.01 | 1 | + |
|  | **Time(Guard)** | 0.73 | 0.25 | 0.26 | 0.71 | 1.27 | 1.01 | 1 | + |
|  | Random effect | Group | Male.id | Oestrus.event |  |  |  |  |  |
|  | sd | 0.324 | 0.782 | 1.16 |  |  |  |  |  |

Numbers (1-4) represent outputs for State transition probabilities (1), Siring probabilities (2), weight loss(3), and mortality proabibility (4) models , with a and b corresponding to age rank (a) and age (b) model variations. Mean (effect), credible intervals (0.025,0.975), and median (50%) effects for each covariate are sampled from the untransformed posterior distribution of each model. f is the proportion of the posterior distribution with the same sign as the mean. Overlap 0 shows whether 0 overlaps with the range of 2.5% and 97.5% quantiles of the posterior distribution for each fitted parameter, and bold covariates are those that had a significant effect (no overlap). Asterisks (*) represent results where posteriors overlapped with 0 but by a marginal amount (f>0.95). Where there was no, or marginal, overlap, the direction of the effect is given in ovelap0. Rhat is a measure of chain convergence (<1.1) [69]. The standard deviation of the three random effects - group, male ID, and oestrus event id - is given for each model.

**Confirmation of a decline in obtaining/maintain a guarding role past the quadratic switch point**

We split the data at the midpoint of the quadratic curve (6 years) and ran separate general linear models for the age effect on young (<=6 years) and old (>6 years) males to obtain and maintain guarding roles. Age had a positive effect on young males to gain a guarding role as opposed to stay as a subordinate (binomial glm: effect=0.688, S.E=0.061, z=11.34, p<0.001) and maintain a guarding tactic rather than drop into a subordinate role (binomial glm: effect=0.560, S.E=0.584, z=9.60, p<0.001). Age had a negative effect on old males to gain a guarding role as opposed to stay as a subordinate (binomial glm: effect=-0.315, S.E=0.125, z=-2.53, p=0.011) and maintain a guarding tactic rather than drop into a subordinate role (binomial glm: effect=-0.362, S.E=0.124, z=-2.91, p=0.004).

**Siring success**

On average 3.83 pups were sired by the group in each oestrus event, not including pups that died before tail tip genetic samples could be taken post litter emergence (aged 30–50 days). The predicted pups sired by an average male was 0.129 (Figure S2a, hci=0.154, lci=0.108). Older age rank and heavier males were significantly more likely to sire a given pup (table 1, model 2a). Age had a significant negative quadratic effect (table 1, 2b), indicative of a decline of the probability of siring at old ages shown past 7 years old (figure S2d).

On average 3.83 pups were sired by the group in each oestrus event, not including pups that died before tail tip genetic samples could be taken post litter emergence (aged 30–50 days). The predicted pups sired by an average male was 0.129 (Figure S2a, hci=0.154, lci=0.108). Older age rank and heavier males were significantly more likely to sire a given pup (table 1, model 2a). Particularly young or light males attained negligible paternity (Figure S2b,c). Males age rank 13 and younger, or 350g or lighter than other males in the group, sired 0.01 or less offspring per oestrus event (figure 4, age rank 13: mean= 0.007, hci= 0.01, lci=0.005; -350g: mean=0.01, hci= 0.015, lci=0.007). Pups sired grew with an increasing gradient as a male moved past age rank 10, or as weight approached the mean (Figure S2b,c). To sire an average of 0.1 offspring per oestrus event, males were predicted to weigh at least the average weight of males in the group (0g: mean=0.01, hci=0.14, lci=0.07), or be of age rank 6 (mean=0.11, hci=0.15, lci=0.08). At the highest age ranks, males were predicted to sire 0.5 or more offspring per oestrus event (age rank 2: mean=0.5, hci=0.65, lci=0.38; age rank1: mean=0.70, hci=0.90, lci=0.53). Extremely heavy males were predicted to sire more pups beyond that predicted by the oldest age ranks, with mongooses weighing at least 400g more than the average male in the group siring at least 1 pup per oestrus event (400g: mean=1.00, hci=1.35, lci=0.76, 450g: mean=1.25, hci=1.64, lci=0.98).

Age had a significant negative quadratic effect (table 1, 2b), indicative of a decline of the probability of siring at old ages shown past 7 years old (figure S2d). Mean pups sired peaked at around 6.5 years (mean=0.81, hci=1.2, lci=0.56), declining by half at 9 years (mean;=0.38, hci=0.60, lci=0.25), and to around 0.1 pups per oestrus event at 10.5 years (mean=0.10, hci=0.17, lci=0.06). Overall, the males with the highest fitness (high weight, old age rank, not over 8 years old) mirrored the males with the highest probability of gaining and keeping a guarding role from event to event.

Figure S2: Predicted mean pups sired per oestrus event given a male’s weight relative to other male group members, age rank, and age. To get an estimate of mean pups sired, the posterior probabilities for siring a given pup were multiplied by the average pups sired in all oestrus events (3.83)

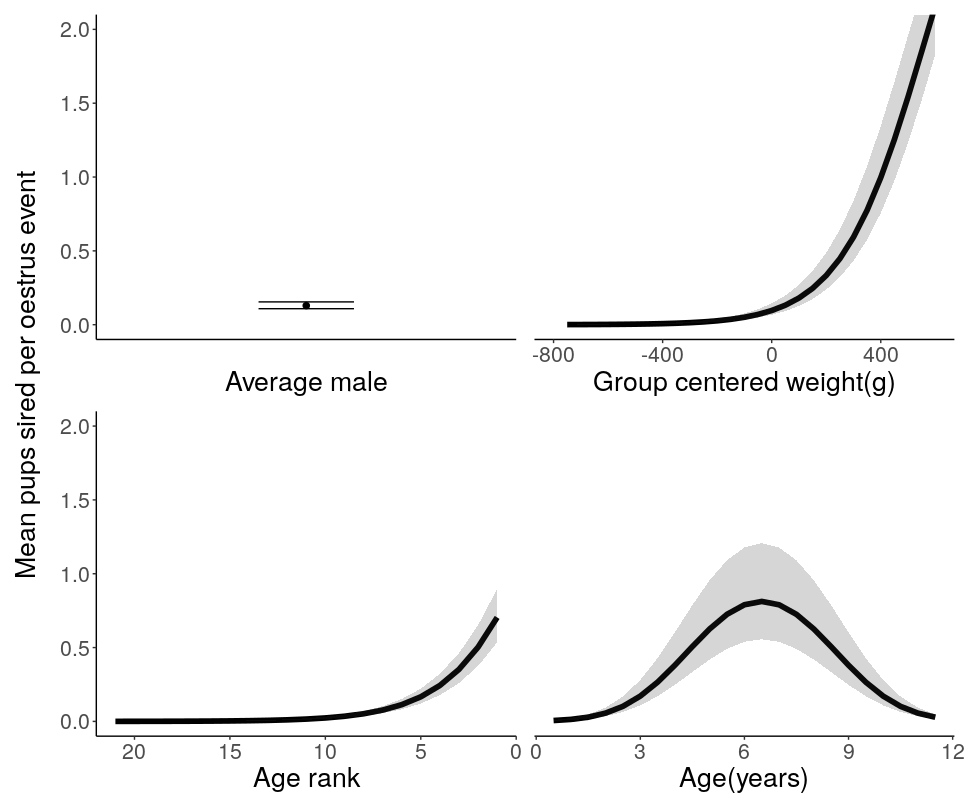

**a	b**

**c	d**

**Weight loss**

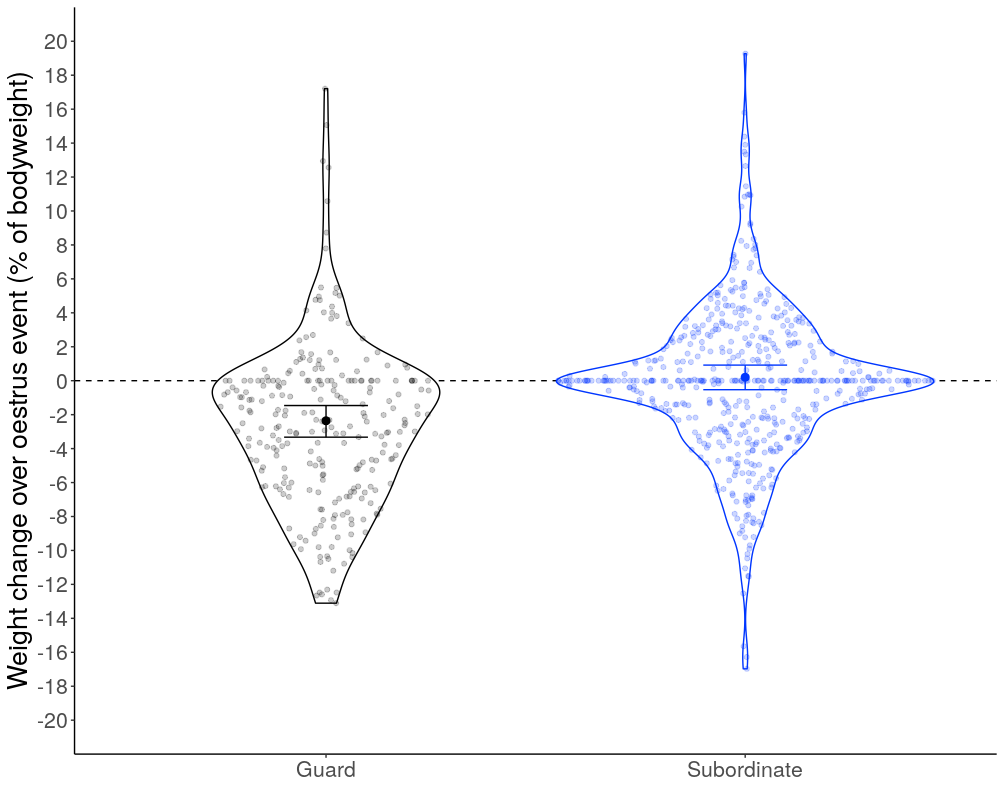

Figure S3: The effect of a male’s reproductive state on their weight change over an oestrus event. Black/grey represents weight change for guarders and blue weight change for subordinates. Single bold points represent the modelled mean weight change, and error bars upper and lower credible intervals. Individual observed weight change is shown by shaded points, and their density distribution represented by a violin plot. The dashed lined represents neutral (0) weight change.
